# Polypolish: short-read polishing of long-read bacterial genome assemblies

**DOI:** 10.1101/2021.10.14.464465

**Authors:** Ryan R. Wick, Kathryn E. Holt

## Abstract

Long-read-only bacterial genome assemblies usually contain residual errors, most commonly homopolymer-length errors. Short-read polishing tools can use short reads to fix these errors, but most rely on short-read alignment which is unreliable in repeat regions. Errors in such regions are therefore challenging to fix and often remain after short-read polishing. Here we introduce Polypolish, a new short-read polisher which uses all-per-read alignments to repair errors in repeat sequences that other polishers cannot. Polypolish performed well in benchmarking tests using both simulated and real reads, and it almost never introduced errors during polishing. The best results were achieved by using Polypolish in combination with other short-read polishers.

**Author summary:** Recent improvements in Oxford Nanopore Technologies sequencing platforms and assembly algorithms have made it easier than ever to generate complete bacterial genome sequences. However, Oxford Nanopore genome sequences suffer from errors that limit their utility in downstream analyses. To fix these errors, one can ‘polish’ the genome with Illumina sequencing, exploiting the fact that Oxford Nanopore and Illumina sequencing have different error profiles. There are several polishing tools which can fix most errors in an Oxford Nanopore genome, but they struggle with errors in repetitive regions of the genome. With this in mind, we have developed a polisher, Polypolish, which uses a novel approach that allows it to fix more errors in genomic repeats. Our results show that Polypolish is both effective at repairing sequence errors and very unlikely to introduce new errors. Polypolish can often fix errors that other polishers cannot and vice versa, so the best results come from using a combination of tools. Polypolish therefore has an important role in bacterial genome assembly methods that aim for the highest possible sequence accuracy.

## Introduction

Long-read-only genome assemblies are inferred using Oxford Nanopore Technologies (ONT) or Pacific Biosciences (PacBio) sequencing reads. For bacterial genomes, reads from these platforms are often longer than the largest repeat in the genome, making complete assemblies (one contig per replicon) possible^1^. However, systematic errors in long reads can lead to hundreds of residual errors in long-read-only assemblies of bacterial genomes, most of which are indels in homopolymer sequences^2,3^. When these errors occur in protein-coding sequences, they cause frameshifts in the open reading frame, leading to problems with genome annotation and limiting the utility of long-read-only assemblies^4^.

Short reads from Illumina platforms do not suffer from the same errors in homopolymer sequences as long reads^5^. Hybrid assembly, using both short and long reads together, can therefore produce sequences which are both complete and highly accurate. Hybrid assembly can be carried out in a short-read-first manner, where long reads are used to scaffold short-read-based contigs into complete genomes, or a long-read-first manner, where short reads are used to correct errors in long-read-based contigs^6^. Given sufficient read depth, long-read-first hybrid assemblies can be more accurate than short-read-first hybrid assemblies, but errors often remain, particularly in repetitive regions of the genome^3^.

Long-read-first hybrid assembly commonly contains three stages: long-read assembly, long-read polishing and short-read polishing^7^. All three stages are important when maximising assembly accuracy. The first stage (long-read assembly) aims to create a complete assembly free of any structural errors, i.e. an assembly where the only errors are small in scale. The second stage (long-read polishing) aims to repair as many of the residual small-scale errors as possible using only long reads, e.g. with a platform-specific tool such as Medaka^8^. This study is concerned with the final stage, short-read polishing, which aims to repair any remaining small-scale errors using short reads.

There are many short-read polishing tools appropriate for bacterial genomes, including HyPo^9^, NextPolish^10^, ntEdit^11^, Pilon^12^, POLCA^13^, Racon^14^ and wtpoa^15^. Except for ntEdit, which uses a *k*-mer-based algorithm, each of these tools relies on short-read alignments. The standard approach to short-read alignment involves placing each read in a single location where it best aligns (with ties broken randomly), and this is how popular aligners such as BWA-MEM^16^ and Bowtie2^17^ behave with default settings. This method is reliable for non-repetitive parts of the genome, but errors in repeat sequences can cause problems. Specifically, if one instance of a repeat sequence contains an error but another instance does not, reads will preferentially align to the error-free instance, leaving no reads aligned over the error (**Figure 1A**). Short-read polishing tools which use the alignments may therefore be unable to fix errors in repeats.

**Figure 1:**
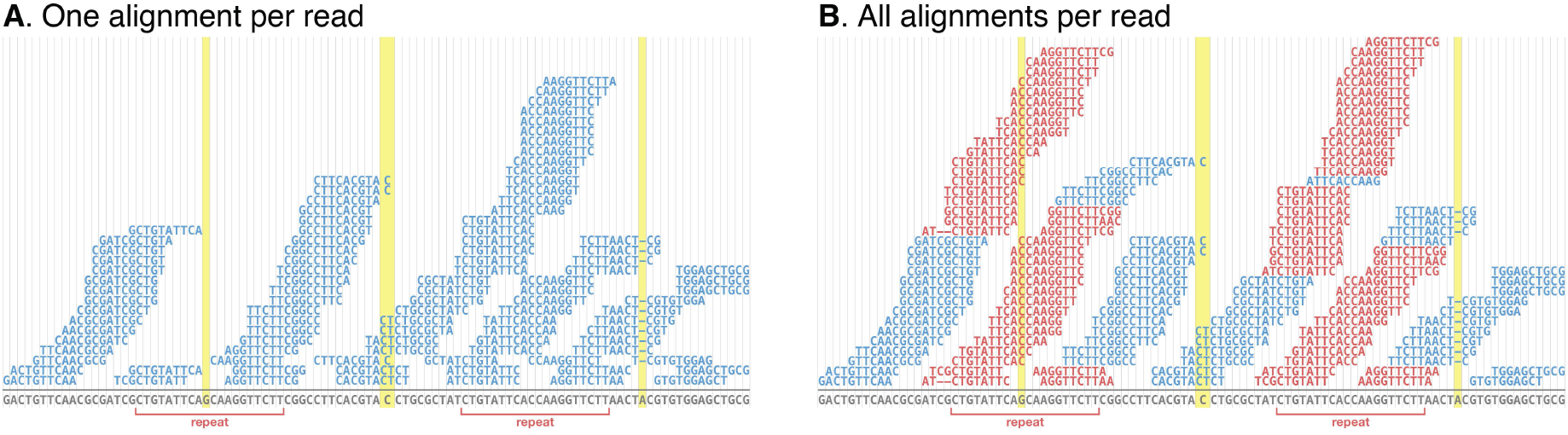
short-read alignments for assembly polishing. In both examples, the aligned reads (above the line) contain no errors and the assembly sequence (below the line) contains three errors indicated by highlighted columns, one of which is in a repeat sequence. **A:** standard one-per-read alignments where each read is aligned to a single best location (randomly chosen in a tie). The errors in non-repeat sequences are covered by read alignments, but the error in the repeat has no coverage because reads preferentially aligned to the other instance of the repeat which is error-free. **B:** all-per-read alignments where each read is aligned to all possible locations. Reads aligned to multiple positions are coloured red. All errors are covered by alignments, including the error in the repeat sequence.

Here, we introduce Polypolish, a new short-read polishing tool which addresses this problem by using a different type of short-read alignment as input. Instead of alignments where each read is placed in a single location, Polypolish is designed to use alignments where each read is aligned to all possible locations (**Figure 1B**). This ensures that errors in repeats are covered by alignments, allowing Polypolish to fix errors that other short-read polishing tools cannot.

## Implementation

Before users run Polypolish, they must align short reads to their long-read assembly using a short-read aligner such as BWA-MEM or Bowtie2. Importantly, the reads must be aligned using the aligner’s all-alignments-per-read option to ensure coverage over genomic repeats (**Figure 1B, Figure S1A**).

Polypolish then builds a pileup, where for each position of the assembly sequence, all the read nucleotides associated with that position are collected (**Figure S1B**). ‘Nucleotides’ in the pileup usually consist of a single base, but a read may contribute more than one base per position in the case of an insertion relative to the assembly. When building the pileup, Polypolish trims off two or more bases at the end of each alignment: whatever base is at the end of the read, however many times it occurs, and one more additional base. This trimming ensures that homopolymer indel errors are properly represented in the pileup (**Figure S2**). Polypolish then calculates a read depth for each position of the assembly (**Figure S1B**). Reads which align to a single location contribute one unit to the depth for each position they cover. Reads which align to multiple locations contribute fractional depth (the reciprocal of their alignment count) to their covered positions; note that this results in approximately the same read depth that would be calculated from the more typical best-match alignment (**Figure S3**).

Polypolish then assesses each base call in the assembly using the pileup and read depth. For any assembly position at which the pileup contains one and only one valid nucleotide, that valid nucleotide differs from the one currently in the assembly, and all other nucleotides in the pileup are invalid, the assembly base call will be changed to the valid nucleotide (**Figure S1C**). Valid nucleotides are defined as those which have a count greater than the valid threshold: 50% of the read depth at that position, or five, whichever is greater (adjustable by the --`fraction_valid` and --`min_depth` options). Invalid nucleotides are defined as those which have a count less than the invalid threshold: 20% of the read depth at that position (adjustable by the --`fraction_invalid` option). In this manner, Polypolish is a conservative polisher, only making a change where the reads indicate a single unambiguous alternative. In all other cases (e.g. when there are multiple valid nucleotides or there are nucleotides between the two thresholds), the assembly base call is left unchanged. Polypolish is therefore very unlikely to introduce errors during polishing.

## Results

### Simulated-read tests

We conducted tests using reads simulated from 100 NCBI reference genomes (details in **Methods, Table S1** to test the effectiveness of Polypolish v0.4.3 and other short-read polishers: HyPo v1.0.3, NextPolish v1.3.1, Pilon v1.24, POLCA v4.0.3, Racon v1.4.21 and wtpoa v2.5. For each genome, we generated a simulated short-read set and an error-containing long-read assembly with an accuracy of ~Q30 (equating to approximately 1000–10000 errors per genome). We then polished the genomes using the short-read polishers and counted the number of remaining errors by aligning the result to the original reference genome (**Table S2**).

We tested each short-read polisher in isolation by running three consecutive rounds of polishing. The resulting per-genome error counts after the third round of polishing are shown in **Figure 2A**. HyPo, NextPolish, Pilon, POLCA and Polypolish performed well, each achieving a mean per-genome error count of <100 and reducing some genomes to zero errors. NextPolish performed best, with a mean per-genome error count of 35. Racon and ntEdit were able to reduce the number of assembly errors, but their mean per-genome error count was >100. wtpoa is excluded from our main results because it increased the number of errors in most genomes (**Figure S4**). Pilon was the tool which most benefitted from multiple rounds (Q46.4 after one round, Q48.8 after two rounds); other tools produced no more than a small increase in accuracy with polishing rounds after the first (**Figure S4**). For most polishing tools, the majority of residual errors were in genomic repeats (**Figure S4**), i.e. the sequence accuracy of non-repeat regions was much higher (>Q50 for most tools) than that of repeat regions (<Q40 for all tools). ntEdit was a notable exception, having a smaller difference between nonrepeat and repeat accuracy (Q40.6 vs Q37.2, respectively).

**Figure 2:**
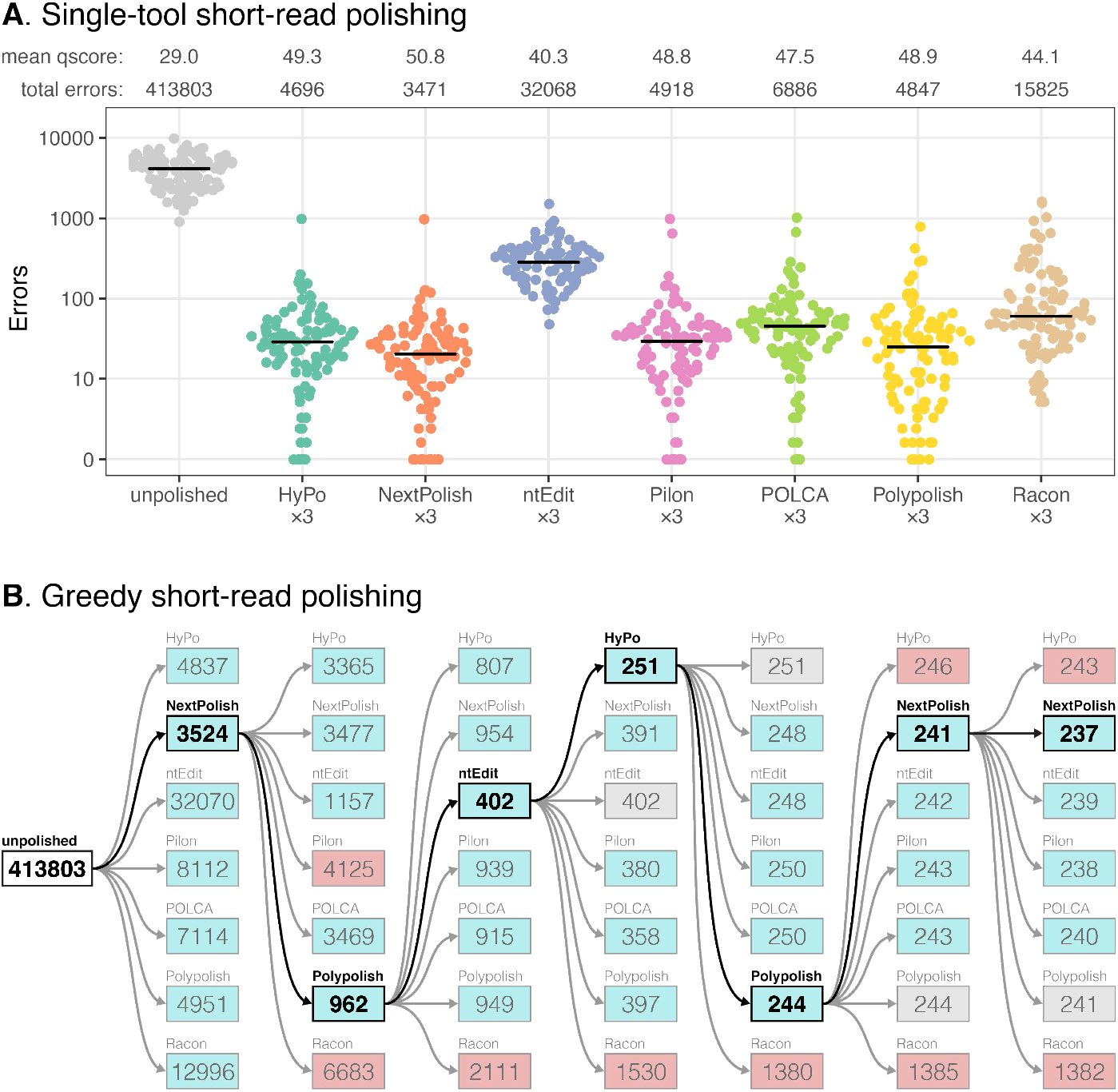
short-read polishing tool benchmarking results using 100 genomes with simulated Illumina reads. **A:** per-genome error rates after polishing with three consecutive rounds with a single tool. Mean qscores and error totals are shown at the top of the plot, and the horizontal lines indicate median error rates for each polisher. **B:** greedy polishing error totals. Each polisher was run on all 100 genomes, and whichever polisher produced the lowest total error count was used as the starting point for another round of polishing with all tools. Decreases in error totals are indicated with a blue box, increases with a red box and no change with a grey box. The lowest total error count for each round is indicated with bold type.

To test ideal combinations of polishing tools, we combined them in an iterative greedy manner. We ran all polishing tools (excluding wtpoa) on all 100 genomes, and whichever output contained the lowest total number of errors was used as the input for the next round of polishing with each tool (**Figure 2B**). This approach achieved 237 total errors across the 100 genomes, less than one-tenth the total error count of any single tool. After seven rounds, 59 genomes had zero remaining errors, but as with single-tool polishing, most remaining errors were in repeat regions (**Figure S5**). To assess the contribution of Polypolish to the greedy combination results, we performed the same approach with Polypolish excluded (**Figure S6**). This resulted in a total error count of 1002 (more than 4× the error count of greedy polishing including Polypolish) and only 31 genomes had zero remaining errors.

To assess each polisher’s likelihood of introducing errors, we quantified false positive changes (per-reference-base introduced errors) that occurred in the first round of polishing (**Figure S7**, **Table S3**). Polypolish achieved the lowest false positive rate (2 introduced errors across all 100 genomes), followed by POLCA (37 errors), HyPo (51 errors), NextPolish (102 errors), ntEdit (371 errors), Pilon (2760 errors), Racon (5161 errors) and wtpoa (8183772 errors). We also ran each polishing tool using the original reference genome as input to test their likelihood of introducing errors into an error-free assembly (**Figure S8**). ntEdit, Pilon, POLCA and Polypolish produced error-free output for all 100 genomes. HyPo introduced an error into three of the genomes, and NextPolish introduced an error into one of the genomes. Racon and wtpoa introduced errors into all 100 genomes. The greedy polishing tests (**Figure 2B** and **Figure S6**) also demonstrate each polisher’s likelihood of introducing errors: HyPo, NextPolish, Pilon and Racon sometimes increased the total error count, while ntEdit, POLCA and Polypolish never increased in total errors.

Hybrid polishing involves using both short and long reads to correct assembly errors. Only two tools, HyPo and Pilon, can perform true hybrid polishing, where both short and long reads are used in the same algorithm (NextPolish, Racon and wtpoa are capable of short-read polishing and long-read polishing but not both at the same time). We therefore tested HyPo and Pilon’s hybrid polishing with the simulated reads to compare their performance to short-read-only polishing with the same tools (**Figure S9**). HyPo-hybrid performed better than HyPo-short (2827 vs 4696 remaining errors across the 100 test genomes after three rounds of polishing), but HyPo-hybrid was much more likely than HyPo-short to introduce errors into an error-free assembly (2550 vs 3 total added errors). Pilon-hybrid performed considerably worse than Pilon-short (231739 vs 4918 remaining errors after three rounds of polishing).

We also used the simulated read sets to explore potential correlates of accuracy that can be calculated without a reference genome and could thus be used to assess genomes assembled using real read sets (**Table S4**). The best overall reference-free predictor of reference-based accuracy was ALE score (**Figure S10**), which is generated by the tool ALE^18^ using short-read alignments to the assembly. ALE scores are calculated using the quality of read alignments, insert sizes inferred from read alignments, evenness of read depth and the assembly’s *k*-mer distribution. ALE scores do not provide an absolute metric of assembly quality but rather a relative metric which can be used to compare alternative assemblies of the same genome, with higher scores suggesting fewer errors.

### Real-read tests

As error-free reference genomes are not available for real read sets, we instead used six isolates for which we had four independent sets of matched Illumina and ONT reads for each isolate (**Figure S11A-B**): *A. baumannii* J9, *C. koseri* MINF_9D, *E. kobei* MSB1_1B, *Haemophilus* M1C132_1, *K. oxytoca* MSB1_2C and *K. variicola* INF345^19^. For each isolate, we produced four long-read-only assemblies (one for each ONT read set) using Trycycler^3^ and Medaka^8^ (**Figure S11C**). We then polished each assembly with an independent set of Illumina data using each of the short-read polishing tools (**Figure S11D**). In the absence of a reference genome against which to directly count assembly errors (as was done for the simulated-read tests), we instead quantified accuracy using ALE scores and the total pairwise chromosomal distance between the four assemblies of each genome that were produced using independent read sets derived from the same isolate (**Figure S11E, Table S5**). Lower pairwise distances imply more accurate assemblies, with a total distance of zero indicating that all four assemblies have identical chromosome sequences.

As with the simulated-read tests, we tested each short-read polisher in isolation by running three consecutive rounds of polishing, as well as combining polishing tools in a greedy fashion (**Table S5)**. Polypolish performed best in the single-polisher tests and was the only polisher which reduced the total pairwise distance to less than 10 for all six isolates (**Figure 3A**, **Figure S12**). Polypolish also achieved the highest mean ALE score for the real-read assemblies. The best results in the greedy combination tests came from one round of Polypolish followed by one round of POLCA (**Figure 3B**). This strategy reduced all genomes to zero total pairwise distance except for one of the four assemblies of *A. baumannii* J9, which had a single difference in a homopolymer in an *ISAba1* sequence (a 6× repeat in this genome)^20^. Greedy polishing without Polypolish was only able to reduce a single genome to zero total pairwise distance (**Figure S13**, **Table S5**).

**Figure 3:**
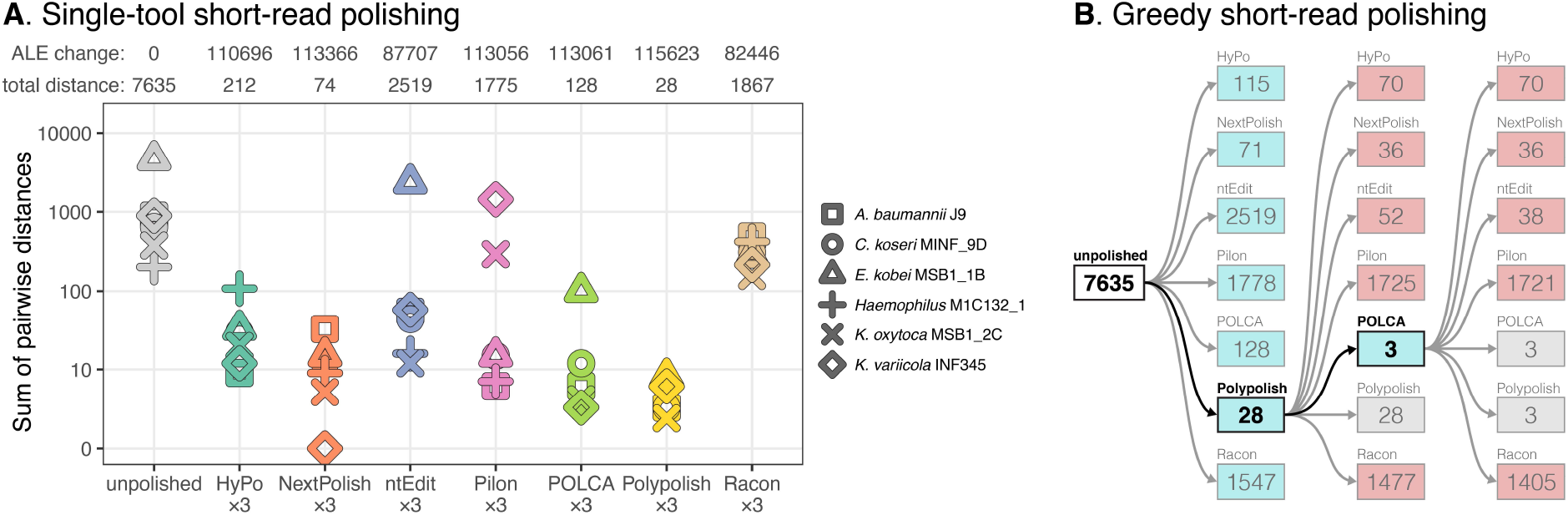
short-read polishing tool benchmarking results using six clusters of genomes with real ONT and Illumina reads. Each cluster contains 3-4 assemblies of the same genome made from independent read sets, each assembled and polished with independent ONT reads and then polished with matched Illumina reads. Instead of quantifying error rates directly (as was done with the simulated-read tests), accuracy is quantified as the sum of pairwise sequence distances within each cluster. **A:** per-cluster distances after polishing with three consecutive rounds with a single tool. Distance totals (lower is better) and mean ALE scores relative to the unpolished genomes (higher is better) are shown at the top of the plot. **B:** greedy polishing distance totals. Each polisher was run on all clusters, and whichever polisher produced the lowest total distance was used as the starting point for another round of polishing. Decreases in distance totals are indicated with a blue box, increases with a red box and no change with a grey box. The lowest total for each round is indicated with bold type.

Some polishers performed poorly on just one or two isolates in the real-read tests, with different isolates yielding the greatest total pairwise distance for different polishers (see **Figure 3A** and detailed descriptions in **Table S5**). Most notably, Pilon introduced large deletion errors into the *K. oxytoca* and *K. variicola* assemblies which did not always result in a decrease in the assembly’s ALE score (**Figure S14**).

## Discussion

By taking all-per-read alignments as input, Polypolish was able to fix errors in long-read assemblies that other short-read polishers could not. This was particularly true for repetitive regions of the genome, where residual errors were most common. While Polypolish did well on its own, the best results came from combining polishers which use different algorithms. For many of our simulated-read tests, a combination of Polypolish with other polishers enabled perfect (zero-error) assemblies (**Figure S5**).

We found that some polishers could introduce errors into assemblies, for example, Pilon created large deletion errors in both *Klebsiella* genomes in our real-read tests. In most real-world scenarios, a ground truth reference genome is not available against which to compare an assembly, so it can be difficult to tell whether a polisher’s changes are fixing or introducing errors. We found that an assembly’s ALE score was a good proxy for its accuracy (**Table S4**, **Figure S10**), so researchers can run ALE before and after polishing, using the difference in ALE score to inform whether any changes are likely to be improvements. However, we did encounter cases where a decrease in assembly accuracy caused an increase in ALE score (**Figure S14**). We therefore recommend using POLCA and Polypolish, the two polishers with the lowest false positive rates (**Figure S7**) and the only two polishers which never made the error/distance totals increase in either our simulated-read or real-read greedy combination tests (**Figure 2B**, **Figure 3B**, **Figure S6** and **Figure S13**).

Since errors in repeats are the most challenging errors to fix in bacterial genomes, there are two strategies which can lead to higher quality assemblies. The first is to improve assembly accuracy before short-read polishing. Long reads are often longer than the repeats in a bacterial genome, so long-read assemblies do not generally yield a pronounced quality discrepancy between repeat and non-repeat sequences. Fewer overall errors in long-read assemblies should therefore mean fewer hard-to-fix errors in repeats. The second strategy is to employ repeat-aware short-read polishing strategies, as is done by Polypolish. We can therefore expect future improvements in long-read sequencing, long-read polishing and short-read polishing to all contribute to making bacterial genome assemblies reliably error-free.

## Methods

### Simulated-read tests

To prepare the reference genomes for the simulated-read tests, we first downloaded all 120 prokaryote reference sequences included in NCBI’s curated subset of high-quality genomes of scientific importance (ftp.ncbi.nlm.nih.gov/genomes/GENOME_REPORTS/prok_reference_genomes.txt). Genomes with multiple chromosomes and those which had problems with long-read assembly (see below) were excluded, leaving 108 genomes. We then identified the eight pairs of genomes that were most closely related (as determined by Mash distance^21^) and excluded one genome from each pair, producing a set of 100 genomes with minimal redundancy (listed in **Table S1**). We excluded plasmids from each genome, leaving only the chromosome, and any ambiguous DNA bases (e.g. N) were replaced with random unambiguous bases (A, C, G or T).

For each genome, we simulated both long and short reads from the reference sequence. Long reads were simulated with Badread v0.2.0^22^ using the following parameters: 100× depth, 90% mean identity, 98% maximum identity, 4% identity stdev, 20 kbp mean read length, 12 kbp read length standard deviation, and all other parameters left at defaults. Paired-end short reads were simulated with ART v2016-06-05^23^ using the following parameters: 100× depth, HiSeqX TruSeq preset, 150 bp read length, 400 bp mean fragment length, 50 bp fragment length standard deviation, and all other parameters left at defaults. All simulated read sets are available in **Supplementary data**.

To identify repeat regions of each reference genome, we simulated deep (300×), error-free, 150 bp, unpaired reads using wgsim^24^. These reads were aligned to their reference genome with BWA-MEM v0.7.17 using the -a option (all alignments per read)^16^. We then used a custom script (find_repetitive_regions.py, available at github.com/rrwick/Polypolish-paper) to parse these alignments and identify all regions of the genome where reads aligned to multiple places. I.e. we defined repetitive regions as those with high enough sequence identity to allow for cross-repeat alignment with BWA-MEM.

To produce an unpolished sequence (an error-containing long-read assembly) for each genome, we first assembled the simulated long-read set using Flye v2.8.3^25^ and then adjusted the strand and starting position of the resulting contig to match the original reference sequence. The errors in the resulting contigs were mostly homopolymer deletions (a consequence of the Badread error profile), so we used a custom script (add_errors.py, available at github.com/rrwick/Polypolish-paper) to add additional error types (homopolymer insertions, non-homopolymer indels and substitutions, each at a rate of 0.01%) to produce the unpolished genomes for the simulated-read tests. These sequences had identities ranging from 99.57% (Q23.7) to 99.94% (Q32.3), and their worst-100-bp identities (the minimum identity in a 100-bp sliding window across an alignment to the reference sequence) were all above 80%, indicating that none of the sequences suffered from large-scale structural errors.

Polished genome sequences were generated by running the short-read polishers using default settings or (if provided) by following recommended commands in their documentation. Exact commands used are available in the wrapper scripts in **Supplementary data**. For the single-tool tests, each polisher was run consecutively three times on each genome, using the polisher’s output assembly as input for the next round. Reference-polishing tests were conducted by running each polishing tool once using the error-free reference sequence as input. For the greedy combination tests, each polisher (excluding the hybrid polishers and wtpoa which performed poorly in the single-tool tests) was run once on each genome, and the best-performing polisher was defined as the one with the fewest total errors in its output assemblies. The best-performing polisher’s output was then used as input for another round of polishing. This process was repeated for a total of seven rounds, after which almost no changes occurred: only four of the 100 genomes changed in the fifth round and only one genome (NC_008596.1) changed in the sixth and seventh rounds. The greedy combination tests were then performed again with Polypolish excluded.

Assembly accuracy was quantified using a custom script (assess_assembly.py, available at github.com/rrwick/Polypolish-paper). This script uses the edlib^26^ library to perform a global alignment of the assembly to the reference sequence. It reports an error count (the number of non-matching positions in the alignment) and identity (the number of matching positions divided by the length of the alignment). It further characterises errors based on their type (substitution, insertion or deletion) and whether they are in a repeat region of the genome. The script then generates other potential metrics of assembly quality using coding sequences and read alignments. It uses Prodigal^27^ to identify potential coding sequences in the assembly (as indel errors are known to cause frame-shift mutations and premature stop codons), reporting the total number of predicted coding sequences, the total length of predicted coding sequences and the mean length of predicted coding sequences. It also uses ALE^18^ to assess the quality of the assembly using short-read alignments (from BWA-MEM^16^), reporting the total mapping quality, total short-read alignment score, ALE score and ALE sub-scores (placement score, insert score, depth score and *k*-mer score).

Confusion matrices (true positive, true negative, false positive and false negative counts across all 100 genomes) were generated for each polisher using a custom script (confusion_matrix.py, available at github.com/rrwick/Polypolish-paper). This script generates a whole genome multiple sequence alignment of the reference sequence, the unpolished sequence and the polished sequence (one polishing round) using Trycycler v0.5.0^3^. It then categorises each position of the genome as a true positive (polisher fixed an error), true negative (polisher did not change a correct base), false negative (polisher failed to correct an error) or false positive (polisher introduced an error) (**Figure S7**).

We assessed potential reference-free metrics of assembly quality using all available data: single-tool results, reference-polishing results and greedy combination results. For each metric, we calculated a Kendall rank correlation coefficient for each genome (true assembly identity as determined by alignment to the reference vs the reference-free metric), and then took the mean of all 100 coefficients (one per genome) to get the metric’s overall correlation coefficient (**Table S4**).

### Real-read tests

The six bacterial isolates used in the real-read tests each belong to a different species: *Acinetobacter baumannii*, *Citrobacter koseri*, *Enterobacter kobei*, an unnamed *Haemophilus* species (given the placeholder name *Haemophilus sp002998595* in GTDB R202^28,29^), *Klebsiella oxytoca* and *Klebsiella variicola.* Sequencing was previously described in Wick et al. (2021)^19^. Briefly, isolates were cultured overnight at 37°C in Luria-Bertani broth and DNA was extracted using GenFind v3 according to the manufacturer’s instructions (Beckman Coulter). The same DNA extract was used to sequence each isolate using three different approaches: ONT ligation, ONT rapid and Illumina (**Figure S11A**). For ONT ligation, we followed the protocol for the SQK-LSK109 ligation sequencing kit and EXP-NBD104 native barcoding expansion (Oxford Nanopore Technologies). For ONT rapid, we followed the protocol for the SQK-RBK004 rapid barcoding kit (Oxford Nanopore Technologies). All ONT libraries were sequenced on MinION R9.4.1 flow cells. ONT read sets were basecalled and demultiplexed using Guppy v5.0.7, using the super-accuracy model. For Illumina, we followed a modified Illumina DNA Prep protocol (catalogue number 20018705), whereby the reaction volumes were quartered to conserve reagents. Illumina libraries were sequenced on the NovaSeq 6000 using SP reagent kits v1.0 (300 cycles, Illumina Inc.), producing 150 bp paired-end reads with a mean insert size of 331 bp. The resulting Illumina read pairs were shuffled and evenly split into two separate read sets, which were combined with the ONT read sets to produce two independent hybrid read sets (**Figure S11B**). We repeated this process (from culture to sequencing) to generate another two hybrid read sets for a total of four hybrid read sets per isolate. All reads are available in **Supplementary data**.

For each hybrid read set, we performed a long-read-only assembly using Trycycler v0.5.0^3^ and Medaka v1.4.3^8^, following the instructions in Trycycler’s documentation (**Figure S11C, Table S6**). One of the ONT read sets for *K. oxytoca* MSB1_2C had very low depth (10×) and was therefore not able to yield a high-quality long-read-only assembly, leaving only three assemblies for this genome. We were able to produce four complete (circularised) long-read-only assemblies for the other five genomes, giving a total of 23 assemblies which served as the ‘unpolished’ assemblies in our real-read tests.

Polished genome sequences were generated by running the short-read polishers as described above (**Figure S11D**). For the single-tool tests, each polisher was run consecutively three times on each assembly. For the greedy combination tests, each polisher (excluding wtpoa which performed poorly in the single-tool tests and the hybrid polishers) was run once on each genome, and the best-performing polisher was defined as the one with the smallest total pairwise distance in its output assemblies. The best-performing polisher’s output was then used as input for another round of polishing until there were no more improvements. The greedy combination tests were then performed again with Polypolish excluded.

To assess the quality of real-read assemblies, we used the edlib^26^ library to perform a global alignment of the chromosome sequences for all pairwise combinations within each genome (**Figure S11E**). The total distance was used as a metric of assembly quality, with lower values being better and a value of zero indicating that all assemblies for the genome were identical. We also ran ALE^18^ on each real-read assembly using short-read alignments from BWA-MEM^16^.

## Supplementary data

The Polypolish tool and documentation can be found at: github.com/rrwick/Polypolish

Supplementary figures, tables, methods and scripts can be found at: github.com/rrwick/Polypolish-paper

Read sets and assemblies can be downloaded from: bridges.monash.edu/articles/dataset/Polypolish_paper_dataset/16727680

